# Optimizing microtubule arrangements for rapid cargo capture

**DOI:** 10.1101/2021.06.02.446824

**Authors:** Saurabh Mogre, Jenna R. Christensen, Samara L. Reck-Peterson, Elena F. Koslover

## Abstract

Cellular functions such as autophagy, cell signaling and vesicular trafficking involve the retrograde transport of motor-driven cargo along microtubules. Typically, newly formed cargo engages in slow diffusive movement from its point of origin before attaching to a microtubule. In some cell types, cargo destined for delivery to the perinuclear region relies on capture at dynein-enriched loading zones located near microtubule plus-ends. Such systems include extended cell regions of neurites and fungal hyphae, where the efficiency of the initial diffusive loading process depends on the axial distribution of microtubule plus-ends relative to the initial cargo position. We use analytic mean first passage time calculations and numerical simulations to model diffusive capture processes in tubular cells, exploring how the spatial arrangement of microtubule plus-ends affects the efficiency of retrograde cargo transport. Our model delineates the key features of optimal microtubule arrangements that minimize mean cargo capture times. Namely, we show that configurations with a single long microtubule and broad distribution of additional microtubule plus-ends allow for efficient capture in a variety of different scenarios for retrograde transport. Live-cell imaging of microtubule plus-ends in *Aspergillus nidulans* hyphae indicates that their distributions exhibit these optimal qualitative features. Our results highlight important coupling effects between microtubule length distribution and retrograde cargo transport, providing guiding principles for the spatial arrangement of microtubules within tubular cell regions.

## I. INTRODUCTION

Microtubules form an essential component of the intracellular transport system, allowing for long-distance distribution and delivery of components driven by kinesin and dynein motors. In eukaryotic cells, microtubules are organized in a wide variety of arrangements depending on cellular geometry and specific biological transport objectives [1, 2]. These architectures range from centrally anchored radial arrays, to swirling or planar-polarized structures nucleated at the cell periphery, to parallel structures in the narrow cylindrical domains of neuronal projections and fungal hyphae [3]. The stark variation in cytoskeletal organization across different cell types raises a fundamental question regarding how the arrangement of microtubules affects cargo transport functionality. Furthermore, pharmacalogical modulation of cytoskeletal architecture by stabilization of dynamic microtubules has been proposed as a potential intervention to reduce transport deficits associated with neurological injury and disease [4].

Many studies have sought to relate the efficiency of cargo transport with cytoskeletal filament arrangements in various contexts. For disordered networks, the dependence of cargo delivery time on filament polarity, bundling, length, orientation, and local density has been established via continuum models and simulations of explicit network architectures [5–10]. Cellular-scale cargo distribution in these models generally relies on multi-modal transport, incorporating processive runs whose direction is determined by the microtubule arrangement, intersperesed with diffusive or subdiffusive motion that allows transition between microtubules [11, 12]. The microtubule architecture thus modulates transport efficiency both by directing processive motion and by determining the rate of capture for cargo in the passive state.

One biologically important objective for intracellular transport is the capture of newly formed cargo and its delivery towards the perinuclear region. Such cargo includes signaling endosomes [13, 14] or autophagosomes [15] formed at distal regions, or COPII-coated vesicles that bud from the endoplasmic reticulum throughout the cell [16]. Because directed motor-driven transport is much faster than diffusion of vesicular organelles, the initial step of cargo capture can play an important role in determining the overall timescale of delivery towards the nucleus.

Cells with long tubular projections, such as neurons and fungal hyphae, provide a particularly convenient model system for retrograde cargo transport. In neuronal axons, microtubules are highly polarized, with their plus-ends pointing towards the distal tip [17]. Similar plus-end-out polarization is observed in the distal segment of multinucleated hyphae for fungi such as *Aspergillus nidulans* and *Ustilago maydis* [18]. Here, we consider the efficiency of cargo capture for transport towards the cell body in these tubular model systems.

Because these geometries are much longer than they are wide, the axial distribution of cargo capture positions becomes particularly important. Given the typical diffusivity of vesicular organelles on the order of *D* ≈ 0.01*μ*m^2^*/*s, it should take on the order of 1 minute for cargo to explore the radial cross-section of an axon or hypha with radius approximately 1*μ*m. By contrast, the time to reach the cell body via pure diffusive transport would range from hours (for a 10*μ*m hyphal tip) to years (in a millimeter-long axon). Cells rely on processive retrograde transport to replace these unreasonably long time-scales with a much more rapid directed velocity on the order of 1*μ*m/s. In hyphal tips, microtubules are nucleated at the final nucleus, so that initial attachment to a microtubule can enable rapid directed delivery to the nuclear region. In axons, the microtubule cytoskeleton consists of filaments much shorter than the axon itself [19]. Although retrograde-moving cargo pauses when reaching a microtubule minus-end, these pauses tend to last only a few seconds, implying that the cargo need not unbind to a diffusive state requiring recapture before proceeding with its directed motion [19]. We therefore focus specifically on the initial capture process of newly created diffusive cargo, as mediated by the axial distribution of capture regions.

The retrograde transport of cargo to the cell body in tubular cells is primarily carried out by cytoplasmic dynein-1 (“dynein” here) motors along polarized microtubules. In some cell types, dynein accumulates near the tips of growing microtubules (plus-ends) and forms enriched pools referred to as ‘comets' that can act as localized capture regions for cargo [20–24]. The placement of comets can be controlled by varying microtubule length (or nucleation sites in axons), and their positioning in relation to where cargo is formed can determine the diffusive search time before initiation of active transport.

We consider the process of cargo binding to a microtubule either at the plus-end or throughout the cytoplasm, and look for possible arrangements of microtubule plus-ends along the axial direction that facilitate this process for various positions of cargo entry. We begin by focusing on simplified microtubule arrangements to capture cargo entering at the distal tip of a cell, loaded preferentially at microtubule plus-ends. Using an analytic one-dimensional model to represent the tubular geometry, we calculate the capture time, defined as the mean first passage time (MFPT) to encounter the capture zone on a microtubule. We derive the conditions for which the MFPT is minimized, and validate the one-dimensional approximation using three-dimensional (3D) Brownian dynamics simulations. The analysis is extended to include broadly-distributed randomized microtubule configurations evaluated for efficiency of cargo capture with different initial distributions and capture-region lengths. General features are established for microtubule arrangements that allow efficient capture for the full variety of explored scenarios. A minimal model of microtubule dynamics highlights how such optimal arrangements may be obtained by tuning microtubule catastrophe rates. Finally, we quantify live-cell images of *A. nidulans* fungal hyphae to demonstrate that observed microtubule distributions in hyphal tips exhibit the general features identified for optimal arrangements.

## II. METHODS

Experimental methods for imaging microtubules in *Aspergillus nidulans* hyphae are provided in the Supplemental Material. The development and implementation of the mathematical model and computational simulations are described below.

### A. Simplified model system for cargo capture

To explore the role of microtubule configurations on cargo capture in a narrow cellular domain, we leverage both an analytically tractable one-dimensional model and three-dimensional Brownian dynamics simulations for cargo motion in a tube.

We consider a tubular domain of length *L* and radius *R*, with *x* = 0 denoting the cell body and *x = L* corresponding to the distal end of the cell (Fig. 1a). We set *R* = 1*μ*m, as appropriate for both neuronal axons [25] and fungal hyphae [26, 27]. The cargo is modeled as a diffusive particle that either enters the cell at the distal tip, or starts uniformly distributed throughout the cell. Cargo diffusivity is set to *D* = 0.01*μ*m^2^*/*s, in accordance with prior measurements for vesicular cargo in fungal hyphae [27]. Microtubules are treated as straight axial filaments, assumed to be scattered uniformly throughout the radial cross-section of the domain. The filaments are polarized, with their minus-ends at the cell body (*x* = 0) and their plus-ends distributed at different axial positions.

**FIG. 1:**
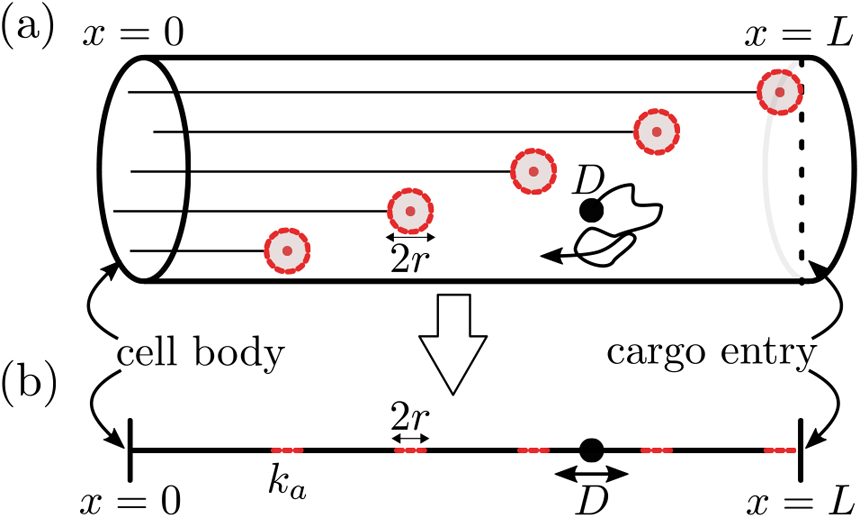
Schematic for cargo capture at microtubule plus-ends in tubular cells. (a) A depiction of a simplified microtubule arrangement in a tubular cell. Cargo enters at *x = L* and diffuses with diffusion coefficient *D*. Cargo is captured at microtubule plus-ends depicted as red circles. (b) Schematic of the equivalent 1D model. A capture rate *k_a_* is introduced to account for the time spent by the cargo diffusing radially at the microtubule plus-end axial location.

**FIG. 2:**
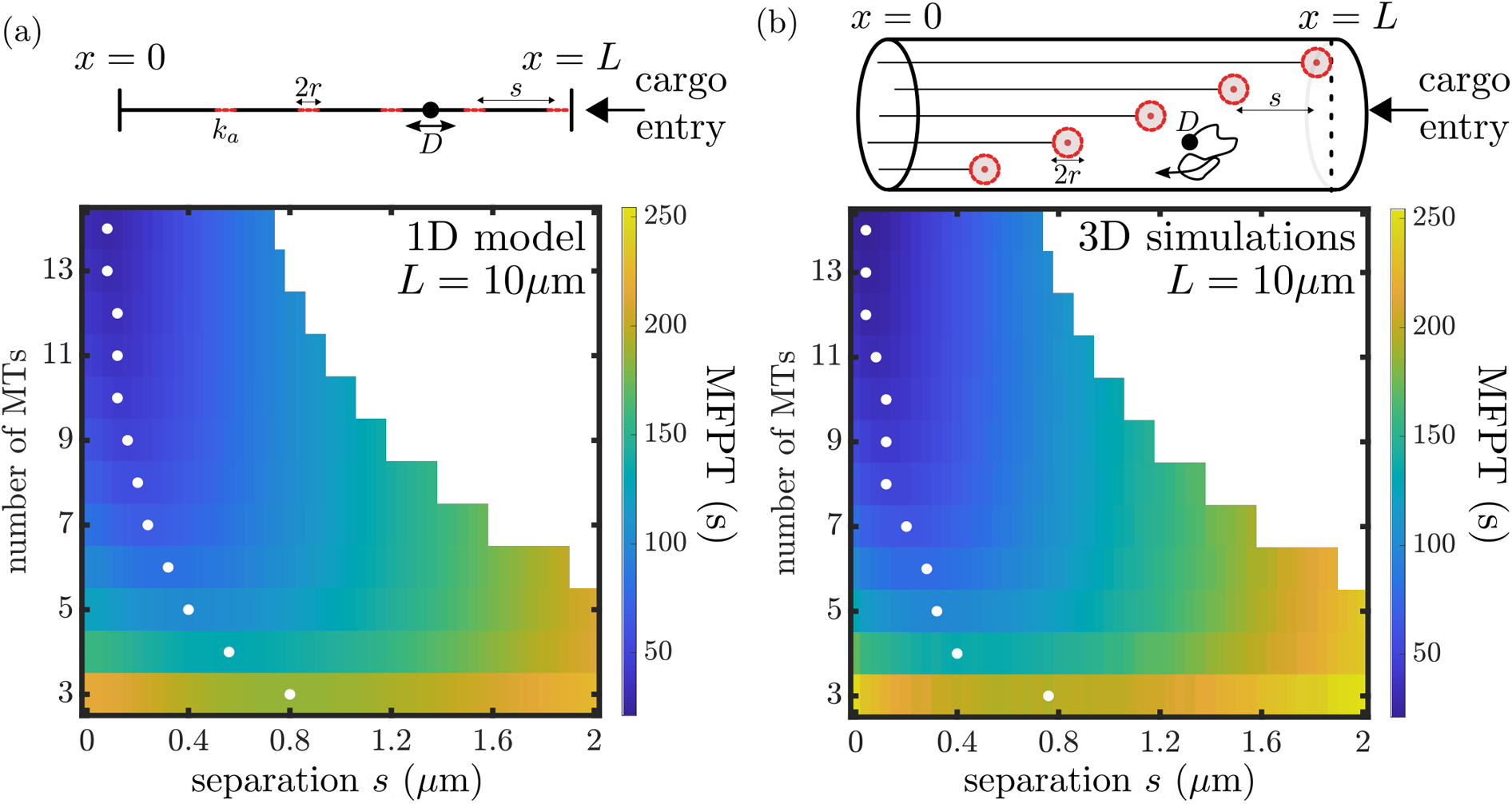
Cargo capture times for axially separated microtubules. (a) Mean first passage time (MFPT) as a function of number of microtubules and separation (*s*) between consecutive microtubules for the effective 1D model. The longest microtubule extends from the cell body at *x* = 0 to the cell tip at *x = L*, which serves as the point of cargo entry. Subsequent microtubules are axially separated by a distance *s*. White dots mark the separation distance *s* that gives the minimum MFPT for a given number of microtubules. (b) Analogous plot for the 3D model of a tubular domain.

**FIG. 3:**
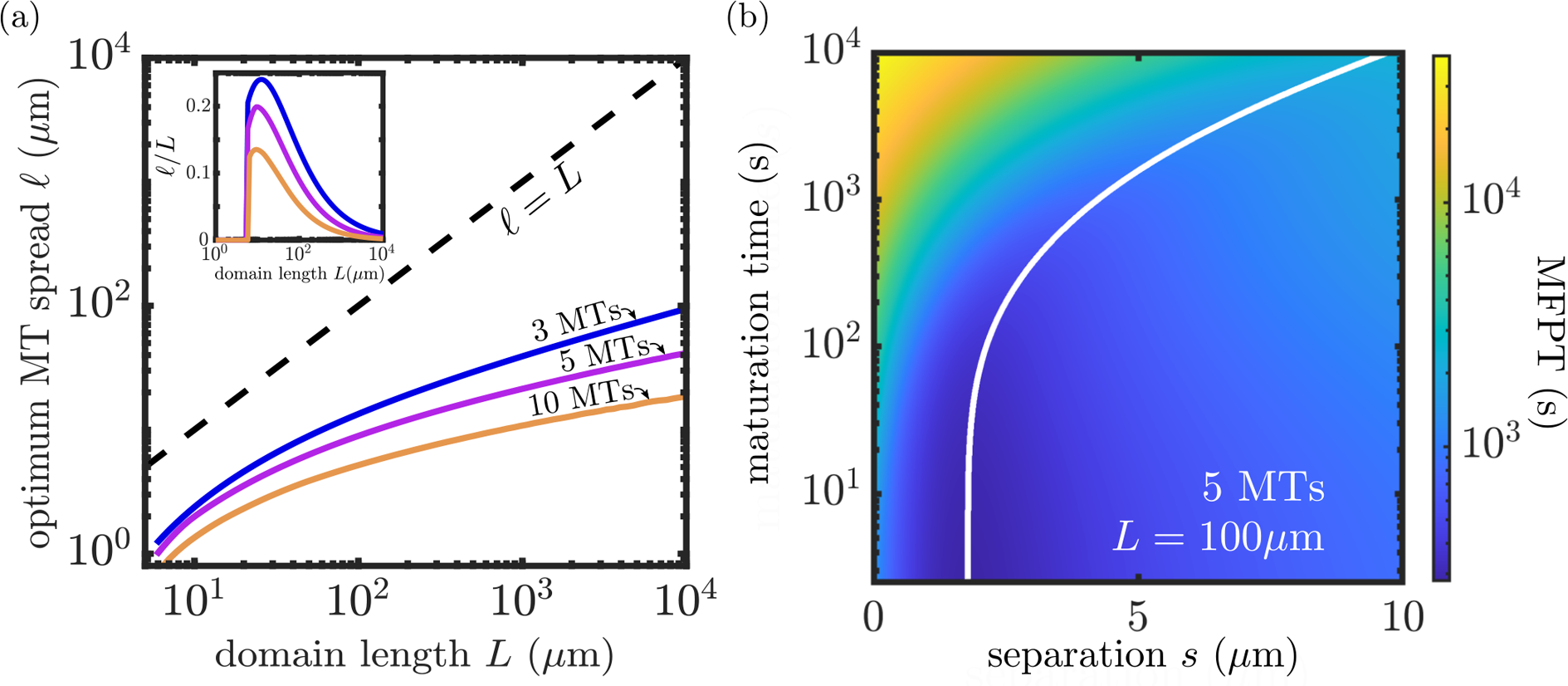
Effect of cell size and cargo maturation on optimal microtubule arrangements. (a) Optimum microtubule spread *l* = *sn_MT_* as a function of domain length *L* for cargos entering at the distal end of the cell and captured at microtubule plus-ends. Inset shows the optimal fraction of the domain length over which microtubules are dispersed. (b) MFPT for maturing cargos entering at the distal end of the cell and loaded at microtubule plus-ends, plotted against the maturation time and separation between consecutive microtubules. The white line denotes the separation with minimum MFPT for a given maturation time. Results are shown for a 100*μ*m cell with 5 microtubules.

**FIG. 4:**
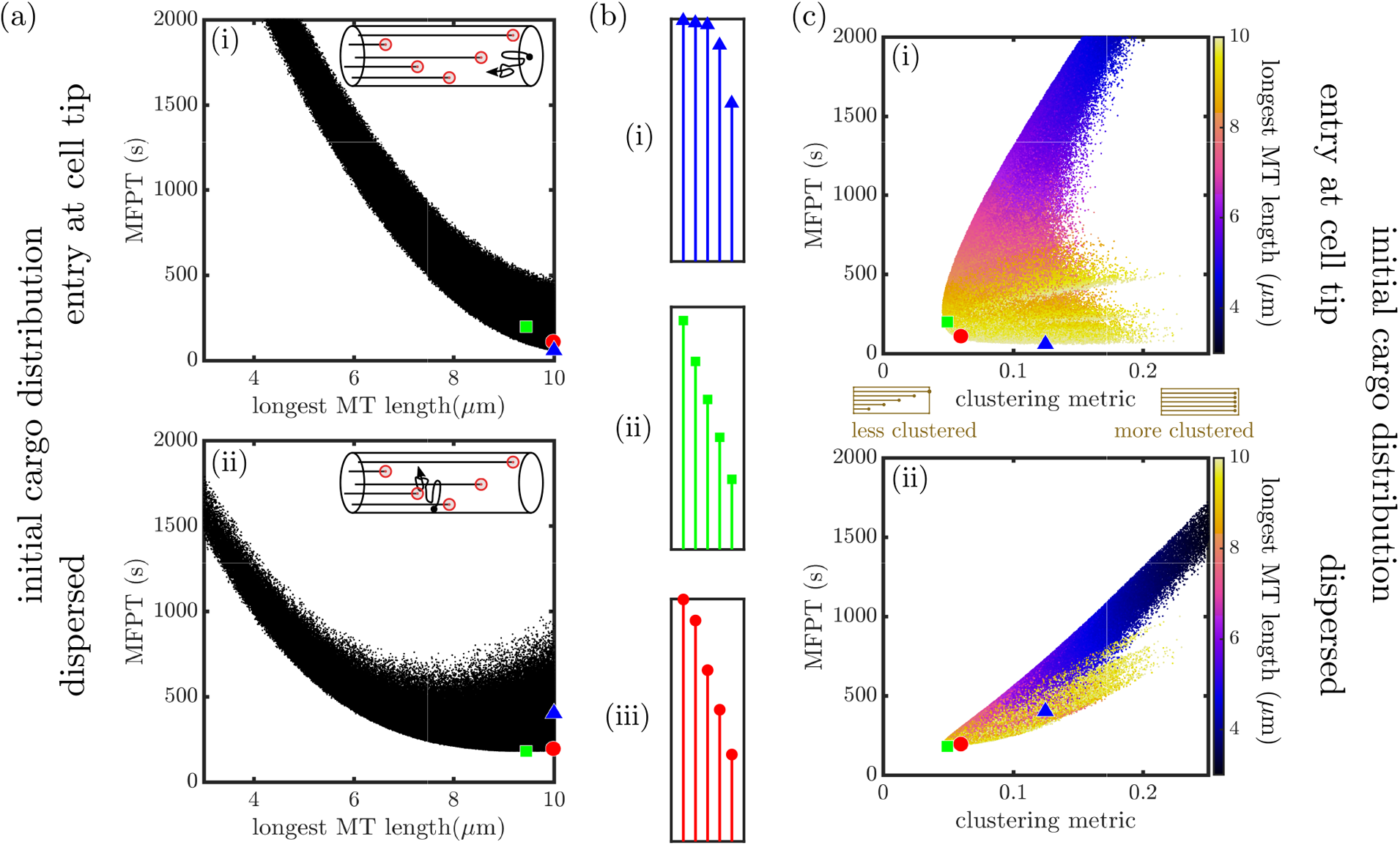
MFPT for random microtubule configurations. (a) Scatter plots showing the MFPT to capture at microtubule plus-ends vs the length of the longest microtubule for 10^6^ randomly sampled configurations with 5 microtubules each in a domain of length 10*μ*m. (i) Cargos start at the cell tip. (ii) Cargos start uniformly. Blue triangle indicates the overall fastest configuration for (i). Green square indicates the overall fastest configuration for (ii). Red circle denotes a configuration that falls within the lowest 2.5% of MFPTs for both starting distributions. (b) Microtubule configurations corresponding to the (i) blue triangle, (ii) green square, and (iii) red circle in panel (a). (c) Scatter plots showing the MFPT plotted against a clustering metric for the randomly sampled configurations, with color indicating longest microtubule length for each configuration. (i) cargos start at cell tip. (ii) cargos start uniformly. Blue triangle, green square, and red circle denote configurations illustrated in panel (b).

Cargo is assumed to be instantly captured when it enters within the capture range (*r* = 0.2*μ*m) of a microtubule plus-end. If the cargo reaches the proximal end of the domain without interacting with a plus-end, it is assumed to have been captured at the cell body.

### B. Analytic one-dimensional model

For very narrow domains (*R ≪ L*), the simple model described above can be mapped to an effectively one-dimensional system, as illustrated in Fig. 1b. The axial positions of cargo and microtubules are projected onto the axis of the cell, represented by a linear segment of length 0 ≤ *x ≤ L*. Cargo can be captured while diffusing within absorbing intervals in the domain, with the rate of absorption determined by the particular arrangement of microtubules.

Plus-ends are denoted as discrete intervals of width 2*r* = 0.4*μ*m, placed at axial positions determined by the microtubule lengths. In an interval corresponding to one microtubule end, the capture rate is set to *k_a_*, representing the rate of encountering the microtubule by diffusion across the radial cross-section. The value of *k_a_* is estimated by computing the mean first passage time in a reflecting cylinder of radius *R* to a central absorbing cylinder of radius *r* [28], according to

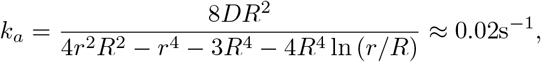

for the estimated parameters *R* ≈ 1*μ*m, *r* ≈ 0.2*μ*m, and *D* ≈ 0.01*μ*m^2^*/*s. For microtubule configurations with axially overlapping plus-ends, the absorption rate is assumed to scale linearly with the number of plus-ends whose capture range overlaps in a given interval.

In this 1D model, a particular arrangement of microtubules can be represented by a series of linear intervals with varying absorption rates. Cargo capture is then represented by a one-dimensional diffusive process in a domain with a reflective boundary at *x = L*, absorptive boundary at *x* = 0, and discrete partially absorbing intervals distributed throughout. The MFPT to capture for this process can be obtained analytically by considering all possible paths of the cargo between the different absorbing segments. Ref. [29] describes a propagator-based approach for computing the mean first passage time to capture a diffusing particle on a network with heterogeneous absorption rates on individual edges. The linear model described here serves as a specialized case of such a network.

For the case of maturing cargo, particles are initialized at the distal end of the domain and are allowed to diffuse freely until they become mature. Only mature particles are capable of capture by the microtubules. The maturation is treated as a Poisson process of rate *λ*. The methodology for incorporating such a process into mean first passage time calculations on a network with heterogeneous absorption rates is detailed in Ref. [29].

### C. 3D simulations for capture dynamics

To validate the approximate one-dimensional model, we also carry out 3D Brownian dynamics simulations of cargo capture by microtubule tips, directly reproducing the cylindrical system illustrated in Fig. 1a. The simulations assume a domain of length *L* = 10*μ*m, and radius *R* = 1*μ*m, reflecting the relevant regime for hyphal tips (region past the last nucleus) in *Aspergillus nidulans* fungal hyphae. Microtubules are modeled as parallel straight lines nucleating at the proximal end of the domain, and are distributed randomly in the radial direction. Diffusing cargos are instantaneously captured when approaching with distance 0.2*μ*m of the microtubule plus-ends.

For a given axial configuration of microtubule ends, the MFPT is computed by averaging over 1000 independent simulations, each sampling a different radial distribution of microtubule positions.

When incorporating microtubule dynamics (in Fig. 5), we turn to a basic model involving microtubule growth and catastrophe. Microtubules are allowed to grow at a speed *v_g_* = 0.18*μ*m/s and shrink at a speed of *v_s_* = 0.62*μ*m/s. These parameters correspond to typical parameters measured in the hyphae of the fungus *Ustilago maydis* [30], which displays similar geometry and transport dynamics to *A. nidulans*. A growing microtubule that reaches the distal tip of the cell remains paused at that location. Both growing and paused microtubules can enter the shrinking state with a catastrophe rate *k*_cat_. A shrinking microtubule switches to the growing state upon reaching the cell body to maintain a constant number of microtubules in the cell. Based on this model, the steady state density of microtubule plus-ends in the cell at an axial position *x* is given by

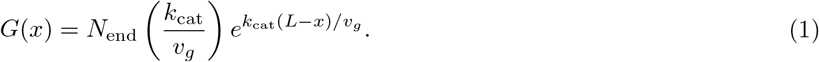

*N*_end_ is the number of microtubules paused at the distal end of the cell, given by

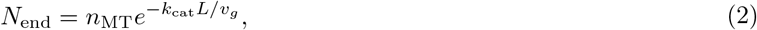

where *n*_MT_ is the number of microtubules in the cell.

**FIG. 5:**
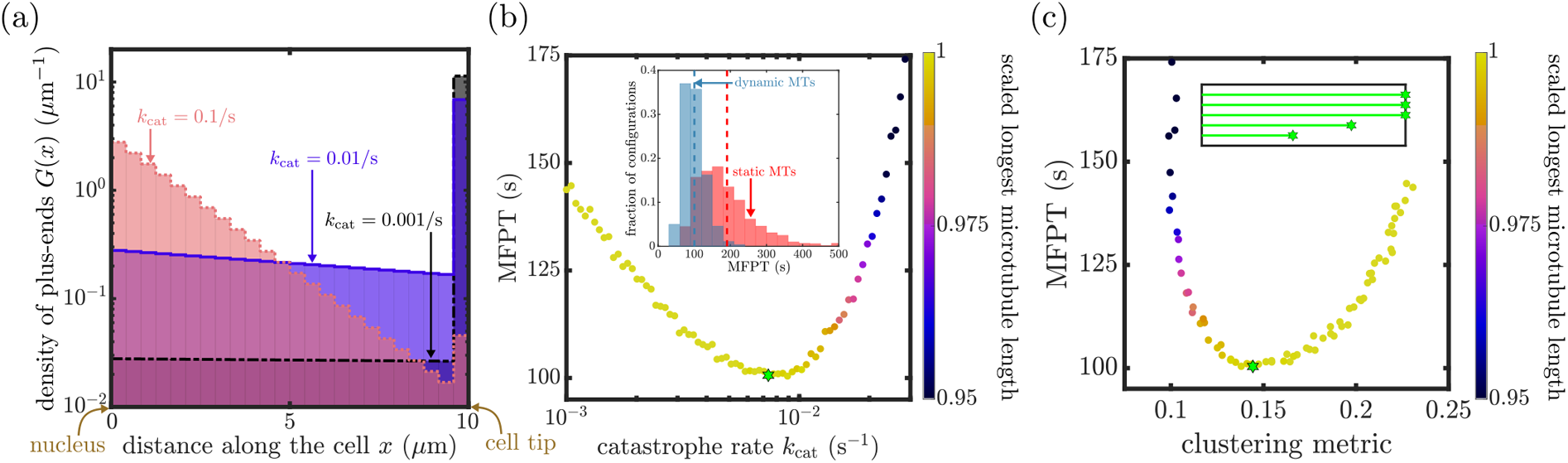
Cargo capture by dynamic microtubules. (a) Steady-state distributions of microtubule plus-ends along the cell axis, for different values of the catastrophe rate (*k*cat). (b) MFPT to capture cargo versus *k*cat for dynamic microtubules. Inset compares the distribution of capture times to simulations with stationary microtubules sampled from the steady-state length distribution corresponding to the optimal value of *k*cat (marked with green star). Dashed lines indicate the mean value for the corresponding distributions. (c) MFPT to capture vs average clustering (minimal-distance metric) for dynamic microtubules. The green star indicates the configuration with minimum MFPT. The inset denotes a representative microtubule configuration corresponding to the optimal catastrophe rate. Color in (b) and (c) indicates the average length of the longest microtubule, scaled by the domain length. All plots are shown for a domain of length *L* = 10*μ*m, radius *r* = 1*μ*m with 5 dynamic microtubules. MFPTs are obtained using 3D simulations, with cargo starting at the distal tip.

Initial microtubule lengths are drawn from this steady state distribution, and microtubules are allowed to grow and shrink according to the described dynamics. Cargo capture at microtubule tips is simulated using the same process as described for static microtubules. We assume that the cargo-capture ability of microtubule plus-ends (*e.g.:* presence of dynein comets) is lost upon catastrophe, so that there is no capture while in the shrinking state.

For a given catastrophe rate, we carry out 1000 independent simulation runs, each starting with a different initial configuration of microtubule ends (uniformly sampled in the radial dimension, and sampled from Eq. 1 and 2 in the axial dimension).

All simulations are carried using custom-built code in Fortran 90, parallelized on the Open Science Grid [31, 32]. Code for both simulations and analytical calculations with the 1D model is provided at https://github.com/lenafabr/transportSimCyl.

### D. Minimal-distance metric to quantify clustering of capture regions

In order to quantify the axial dispersion or clustering of capture regions (*i.e.:* microtubule plus-ends), we define a “minimal-distance” metric for a configuration of *n*_MT_ points on a unit interval. Namely, this metric measures the average distance between a uniformly distributed probe and its nearest point in the configuration.

The configuration is described by normalized points 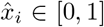, with 1 ≤ *i ≤ n*_MT_. An additional point *x*_0_ = 0 is included to represent absorption at the cell body. Mid-points between consecutive absorbing points are then given by 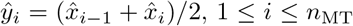. End-points of the domain are denoted by 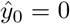 and 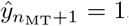, respectively. The closest point for a random number *u* distributed uniformly between 0 and 1 is located at 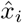 if 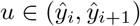. The average distance between a uniformly distributed probe and its nearest absorbing point is then given by,

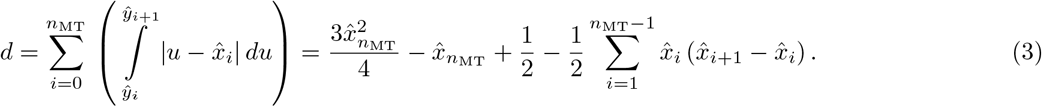

We use this average minimal-distance *d* as the clustering metric throughout the text. Smaller values of *d* correspond to well-dispersed microtubule plus-ends, with a minimal value of 1*/*(4*n*_MT_ + 2) for the configuration where consecutive points are equally spaced. Larger values indicate clustering of microtubule plus-ends along the axis, with a maximal value of *d* = 0.25 for the configuration with all plus-ends at the distal tip.

## III. RESULTS AND DISCUSSION

### A. Separation of microtubule ends for distal capture

We first consider the problem of optimizing the spatial distribution of a limited number of capture regions (*e.g.:* dynein comets at microtubule plus-ends) for rapid capture of diffusive cargo entering at the distal tip of a tubular cell. Given the long lengths (10 – 100*μ*m) and narrow widths (~ 1*μ*m radius) for cellular projections such as fungal hyphae and neuronal axons, the distribution of plus-ends along the axial direction is expected to have the greatest effect on capture times. For simplicity, we assume that captured cargo is rapidly transported directly to the cell body, so that we focus on the rate of the initial capture process.

To begin with, we consider two extreme arrangements of microtubule plus-ends in a long tubular cell. On the one hand, clustering plus-ends near the distal tip will enable distally-produced cargos to quickly encounter and bind to the microtubule. However, any cargo that diffuses past the clustered plus-ends may then embark on very long trajectories down the tube, resulting in a long-tailed distribution of capture times. In general, a diffusive particle that starts at distance *x*_0_ from one absorbing end of a domain of length *L* will have a mean first passage time of *τ = x*_0_(*L − x*_0_)*/*(2*D*) to reach the ends, a quantity that approaches infinity as the domain becomes infinitely long. By contrast, scattering capture regions broadly throughout the domain ensures a uniform availability of capture regions and precludes very long trajectories prior to capture. However, if the number of microtubule tips is fixed, such a broad distribution results in a lower density near the distal origin of the particles and forces each one to diffuse further along the axis before encountering a tip. To quantify this trade-off, we compute the mean first passage time (MFPT) to capture distally-produced diffusing cargo for different spacings of microtubule ends away from the distal tip of the cell.

Specifically, we focus on regularly spaced configurations to explore two key parameters that play a role in cargo capture (Fig. 2a). First, the number of microtubules (*n*_MT_) determines the number of capture regions that a cargo can attach to, with a higher quantity generally corresponding to faster capture. Second, the axial separation (*s*) of consecutive microtubule ends tunes the breadth of their distribution away from the point of cargo entry.

An important advantage of a narrow tubular geometry is that it can be modeled analytically as an approximately one-dimensional system. Since the length of our domain is typically much larger than the radius, a simplified model that represents the tube as a line with localized binding regions can encompass the overall behavior of the capture process. The one-dimensional approximation (Fig. 2a, top panel) represents the microtubule tips as short intervals with a finite capture rate within each interval that encompasses the rate of radially encountering the microtubule while within that slice of the domain.

The geometric parameters of the model reflect a typical hypha of the fungus *Aspergillus nidulans*, which serves as a convenient model system owing to its neuron-like geometry and genetic tractability. We model the distal segment of the hypha (beyond the last nucleus) as a one-dimensional interval with an absorbing boundary at *x* = 0 (corresponding to the cell body) and a reflecting boundary at *L* = 10*μ*m (corresponding to the distal tip) that serves as the original cargo entry point. Microtubule plus-ends are treated as capture regions of width 2*r* = 0.4*μ*m, representing a reasonable capture radius for an organelle and a dynein comet. The first capture region is located at the distal end of the domain, and subsequent regions are placed with a separation *s* from the previous. Once the cargo is within a capture region, it can be absorbed at a rate *k_a_*, representing loading at a plus-end. The capture rate accounts for the time spent by the cargo diffusing radially at the microtubule plus-end location. The average time to capture includes trajectories that pass through multiple capture regions until successfully undergoing absorption within one of them. The MFPT of this process can be computed using a previously developed method for reaction rates on heterogeneous tubular networks [29], as described in Materials and Methods. Fig. 2a shows a plot of the MFPT versus the number of microtubules and the separation between consecutive microtubule ends.

To verify the validity of the approximate one-dimensional model, we compare our results to 3D Brownian dynamics simulations, that encompass the full model system with a tubular domain and spherical capture regions of radius *r* representing the microtubule ends. Details of the 3D model are provided in the Methods section. As shown in Fig. 2, the 1D analytic calculations and 3D simulations give nearly identical results for the MFPT. The mean relative error in the MFPT between the 1D and 3D approaches is ~6.6%. This close correspondence establishes the robustness of the approximate 1D model for representing the narrow tubular geometry. We proceed to employ the 1D model for the remainder of the calculations discussed below.

Interestingly, the MFPT to capture shows non-monotonic behavior as separation *s* is increased from 0, reaching a minimum at an intermediate separation distance between microtubule ends. The existence of the minimum is a consequence of the competition between capturing cargos quickly near the entry point, and extending the overall capture region for cargos that might evade the initial cluster. As the number of microtubules, viz. the overall capture capacity, is increased, the optimal separation decreases. This follows from the fact that fewer cargos can escape the initial capture near the tip when a large number of microtubules are present. The optimal separation ranges between ~0.01*μ*m for 14 microtubules to ~0.8*μ*m for 3 microtubules. Converting the optimal separation to the overall distance over which plus-ends are scattered, the results indicate that it is optimal to distribute plus-ends over a distance of ~1.4 − 2*μ*m from the cell tip for a 10*μ*m cell.

The existence of an optimum separation distance for a given number of microtubules highlights the benefit of scattering capture sites for cargo generated at the cell tip. Intuitively, scattered configurations of microtubule ends are more effective in that they are able to capture cargo that diffuses past the distal region, precluding very long trajectories that explore a large fraction of the domain prior to returning for capture. Because the MFPT between two absorbing boundaries scales in proportion to the domain length, we would thus expect the optimal microtubule end separation to be larger for longer domains. We therefore proceed to explore the effect of domain length on the optimal microtubule distribution.

### B. Effect of cell length and cargo maturation

While many tubular cell projections exhibit a similar width of 1 − 2*μ*m, the length from the distal tip to the nearest nuclear region can vary widely. In fungal hyphae, the distance from the hyphal tip to the first nucleus can range from a few to tens of microns in length [26, 27]. Neuronal axon lengths can vary from hundreds of micrometers up to a meter long [33]. In Fig. 3a, we compute the optimal separation between microtubule ends for cylindrical domains of different length, where the domain length represents the axial distance from the tip to the nearest nucleus.

As expected, the optimum separation increases for longer domain lengths. However, the rate of increase is distinctly sublinear with *L*. This effect arises because as the region containing the microtubule ends becomes longer, it is increasingly likely that the cargo is captured before leaving to explore the rest of the domain. Because this initial capture process is independent of the domain length, the dependence on *L* becomes increasingly shallower as the microtubule ends are more spread out. Consequently, for long tip-to-nucleus distances, an optimum arrangement of microtubule ends concentrates them over a small fraction of this distance. Cell projections of length 10 − 20*μ*m require the widest relative separation of capture regions (Fig. 3a, inset), scattering the microtubule ends over ~ 15 – 25% of the domain.

The scattering of microtubule ends engenders a trade-off between rapidly capturing cargo at its point of entry, and minimizing search time for cargo that wanders within the cell. The optimum distribution therefore depends on the location within the cell where cargo first becomes capable of interacting with the microtubule plus-ends. In some cases, the newly formed cargo may not be immediately available for capture, requiring additional maturation steps such as the acquisition of various adaptor proteins. One such example involves autophagosomes in neuronal axons, which are thought to require lysosome fusion events prior to engaging in retrograde motility [34–36]. We analyze optimal microtubule distribution for capture of cargo with a finite maturation time during which it moves diffusively without being able to bind to microtubule ends.

Varying the maturation rate effectively results in tuning the initial distribution of capture-ready cargo. For example, a very slow maturation rate results in a nearly uniform distribution of cargo available for capture since there is more time to diffuse before maturing. On the other hand, instantaneous maturation reverts to the previously studied case of capture-ready cargo entering at the cell tip. Maturation of cargos can be incorporated as a stochastic Poisson process in the cargo capture model (see Methods).

Fig. 3b shows the MFPT plotted against the maturation time and the separation between consecutive microtubules. As before, there is an optimum separation that minimizes the MFPT for each maturation time. The optimum separation increases as cargo maturation slows down, highlighting the need to spread microtubule plus-ends further in order to capture diffusively wandering cargo that matures slowly. For a 100*μ*m cell, the optimal separation of microtubule tips begins to increase only for maturation times above 100 s.

The dependence of optimum separation on cargo maturation underscores the effect of initial cargo distribution on the MFPT. While there is still a trade-off between clustered and dispersed microtubule plus-ends, matching the location of capture regions to the starting distribution of the cargo leads to more efficient capture. Indeed, a more general treatment would account for various initial cargo distributions. These can be incorporated in the model as initial conditions ranging between the extremes of cargo entering at the cell tip, and cargo being distributed uniformly within the cell.

### C. Optimal microtubule configurations for multiple capture conditions

In the previous section, we focused on cargo produced at the distal tip, and loaded onto microtubules only within a 200nm contact radius of the plus-end. However, both of these assumptions do not necessarily hold for all retrograde transport systems. For example, while the distal tips of hyphae are the most endocytically active [37], some endosomes may be produced elsewhere along the membrane. Other organelles, such as peroxisomes, may bud from the endoplasmic reticulum all along the hyphal length. We therefore consider for comparison the extreme case of cargo produced uniformly throughout the extended cell region. Furthermore, dynein comets exhibit a gradual decrease in density over a micrometer length scale [21], so that capture may not be limited to such a short range of the microtubule plus-end. In this section, we explore the overall features of optimal microtubule configurations for retrograde transport initiation in a variety of cargo production and capture conditions.

We generate 10^6^ random configurations of 5 microtubules, with plus-end positions selected uniformly at random across the domain. For each configuration, we compute the MFPT for capture at the microtubule plus-ends both for distally-initiated and uniformly-initiated cargo. In the Supplemental Material, we also provide results for plus-end capture regions of increasing length, including the extreme case of capture everywhere along the microtubule.

One of the key features of the microtubule configuration is the extent to which it covers the entire cellular domain. This is particularly important for the case of cargo entry at the distal tip, where the presence of a microtubule end near the entry point can greatly speed up capture. We use the length of the longest microtubule in each configuration to describe this feature, demonstrating that the capture time generally decreases as the longest microtubule length is increased (Fig. 4a.i). The optimal configuration in this case involves half the microtubules at near-maximal length, with the remainder scattered up to 3*μ*m away from the tip (Fig. 4b.i). It should be noted that for the case of very long capture regions, the dependence on the longest microtubule length becomes an even stronger predictor of the capture time, with little variability among different configurations that have the same longest length (see Supplemental Material).

When cargo is produced uniformly throughout the domain, the capture efficiency is not so well-correlated with the length of the longest microtubule (Fig. 4a.ii). In this scenario, the optimal microtubule architecture exhibits a broad dispersion of the plus ends throughout the domain (Fig. 4b.ii), in keeping with the broad initial distribution of the cargo. We threfore sought to establish another metric that quantifies the extent to which plus-ends are dispersed or clustered throughout the domain.

For cargo produced at the distal tip of the cell, the length of the longest microtubule determines the position of the nearest capture region relative to the point of cargo entry. For cargo initiated uniformly across all axial positions, we can define an analogous quantity which we term the minimal-distance clustering metric (*d*). Namely, for a given set of microtubule end positions, we compute the expected value of the distance between a point selected uniformly at random and the nearest microtubule end to that point (see details in Supplemental Material). Because particles are also captured at the soma, a capture region at *x* = 0 is appended to all microtubule configurations. The minimal distance metric measures the clustering of capture regions: high values correspond to highly clustered microtubule plus-ends (with a maximal value of *d*_max_ = 0.25); low values correspond to plus-ends spread evenly out over the entire domain (minimal value *d*_min_ = 1*/*(4*n*_MT_ + 2)).

The MFPT to capture at microtubule ends is plotted versus this clustering metric for each of the sampled microtubule configurations in Fig. 4c. When particles start at the distal end of the domain, the capture times are largely insensitive to the clustering metric (Fig. 4c.i). The most optimal (lowest MFPT) configuration for cargo produced at the distal tip (blue triangle) has a moderately high clustering metric (*d* = 0.12), corresponding to slightly separated ends near the distal tip (see Fig. 4b.i), similar to the optimum found in Fig. 2.

By contrast, when cargo starts uniformly throughout the domain, lower clustering ensures that there is always a capture region close to the starting position of the particle, allowing for faster capture times (Fig. 4c.ii). The optimal configuration sampled for this scenario (green square, illustrated in Fig. 4b.ii) has a relatively low clustering metric of *d* = 0.049, close to the minimal possible value of this metric (*d*_min_ = 0.046 for *n*_MT_ = 5). This effect arises because clustered configurations near the cell tip require the dispersed cargo to diffuse over long distances through the cell before it can either reach the cell body or the plus-ends located near the cell tip. On the other hand, evenly dispersed plus-ends provide capture regions throughout the cell, so that all cargos have a capture region nearby regardless of where they initiate.

Our analysis shows that the length of the longest microtubule is a strong predictor of the MFPT when cargos are are captured by long regions of the microtubule (see Supplemental Material), and a moderate predictor when distally produced cargos are captured at the microtubule ends (Fig. 4a.i). In the case of uniformly dispersed cargo captured at microtubule ends, the minimal-distance clustering metric is a complementary predictor of capture efficiency (Fig. 4c.ii). Microtubule configurations that fulfill both of these criteria (high longest length and low clustering) are expected to yield a fast capture time for all of the scenarios considered. We identify a set of 14 microtubule configurations that fall in the lowest 2.5% of MFPT for both distally produced and uniformly produced cargos. Of these, the configuration with the lowest MFPT for distally produced cargos is shown with red circles in Fig. 4. The configuration has one microtubule reaching nearly to the end of the domain and the other microtubules distributed roughly evenly over more than half of the domain length (Fig. 4b.iii). Such a microtubule architecture also exhibits near-optimal capture times (within the lowest 3%) for the extreme case of capture everywhere along the microtubule (Supplemental Material).

Overall, these findings highlight the features of ideal microtubule configurations for efficiently capturing cargo for retrograde transport. Namely, configurations with one long microtubule and other microtubules of broadly distributed lengths result in near-optimal capture times regardless of whether cargos are produced distally or throughout the domain, and of whether they are captured by point-like dynein comets at microtubule ends or more broadly along the whole microtubule.

### D. Establishing optimal configurations through microtubule dynamics

The results above demonstrate the overarching features of microtubule configurations that result in optimal cargo capture. Our model is agnostic as to the dynamic processes by which a cell might establish such an optimum configuration. Furthermore, a key simplifying assumption of the model is that individual microtubule architectures remain fixed throughout the capture process, so that the distribution of microtubule lengths serves as a source of quenched disorder for the position of the capture regions. Realistically, microtubules in fungal hyphae grow and shrink on roughly 30sec timescales [30]. Microtubules in growing neuronal projections are similarly dynamic, although those in mature axons tend to remain relatively stable over time [38, 39].

We briefly consider the effect of microtubule dynamics on the capture process by incorporating basic growth and catastrophe processes into the 3D simulation, as described in Materials and Methods, while fixing a total number of *n*_MT_ = 5 microtubules. The minimal dynamic microtubule model fixes growth and shrinkage velocities according to published data in fungal hyphae [30]. The catastrophe rate *k*_cat_ sets a time-scale on which a growing microtubule halts and begins to shrink, and is used as a free control parameter to tune microtubule distributions. Microtubules that reach the end of the domain are assumed to be capped and to remain fixed until a catastrophe event occurs.

The catastrophe rate modulates the steady-state distribution of microtubule lengths (Fig. 5a). In this simple model, the length of the longest microtubule in the domain and the clustering of microtubule ends are coupled together. Low values of *k*_cat_ result in most of the microtubule plus-ends accumulating at the distal tip of the domain, corresponding to a high value for the longest MT length and for the clustering metric. Intermediate values of *k*_cat_ allow the microtubule ends to spread more broadly through the domain, while high values result in substantial shortening of all microtubules.

We carry out simulations with dynamic microtubules, focusing on the mean time to capture by microtubule plus-ends for particles starting at the distal tip of the cell. Due to the coupling between the longest microtubule length and the end clustering, an optimum value of *k*_cat_ ≈ 0.007s^−1^ emerges for minimizing capture time (Fig. 5b). For this value, the longest microtubules are still able to reach the distal tip of the cell, but other microtubule ends remain relatively well scattered over a broad span of the distal region, as indicated by an average clustering metric of *d* ≈ 0.14 (Fig. 5c). This optimal catastrophe rate is within the range of the measured values (0.006 – 0.04s^−1^) in a variety of cellular systems [30].

We note that the absolute values for the capture times are substantially lower when microtubule dynamics are included in the simulation (Fig.5b, inset). This difference arises from a combination of two effects. First, growing microtubule ends sweep through the domain, tending to pick up any particles that have meandered away from the distal region. Second, the ability of dynamic microtubules to sample several configurations over the hundred-second timescale of particle capture makes it more likely that some microtubule end will encounter the particle, precluding the occasional very long trajectories associated with particles having to return to the distal end for capture. These results highlight a potentially important role for microtubule dynamics in the capture of cargo for retrograde transport.

Despite the overall faster capture times, the dynamic model reproduces the overall features of optimal microtubule end configurations for particle capture. The existence of an optimal catastrophe rate further highlights the balance between allowing a few microtubules to stretch to the distal end of the cell while retaining a broad distribution of microtubule ends throughout the domain.

### E. Microtubule arrangements in *Aspergillus nidulans*

The theoretical work described here provides guiding principles for the performance of different microtubule architectures in capturing cargo. A logical avenue for further study would be to quantify microtubule configurations in actual cellular domains, and to compare the distributions observed with the features identified for optimal capture. To this end, we image hyphae of the fungus *Aspergillus nidulans* and visualize microtubule plus-ends along the hyphal axis.

*A. nidulans* is a filamentous fungus that forms multinuclear tubular projections (hyphae). Owing to its genetic tractability and simplified geometry, *A. nidulans* has been used as a model organism for studies of microtubule based transport [18, 26, 40]. In the hyphal region beyond the most distal nucleus, microtubules form parallel, polarized arrangements, with plus-ends growing towards the distal tip [18]. The distal hyphal segment is on the order of 10*μ*m in length and 1*μ*m in radius [41, 42], allowing it to be approximated as a narrow, effectively one-dimensional tubular region. Endosomes carrying signaling particles are thought to initiate primarily at the distal tip [43, 44], while other organelles, such as peroxisomes, may form by fission or budding from the endoplasmic reticulum throughout the hyphal axis [45].

*A. nidulans* germlings (spores that have recently germinated to form hyphae) expressing GFP-tagged microtubules (tubulin TubA-GFP), mCherry-tagged microtubule plus-ends (microtubule plus-end associated protein EB1 [EbA]-mCherry), and mCherry-tagged nuclei (histone H1 [HH1]-mCherry) were imaged using spinning disk confocal microscopy (details in the Supplemental Material). Sections of the hypha extending from the last nucleus to the cell tip were chosen for analysis. Hyphal length from nucleus to tip was determined by tracing a line along the axis from the end of the last nucleus to the cell tip (Fig. 6a, left). Microtubule plus-ends were enumerated by counting EbA-mCherry puncta within the region beyond the last nucleus (Fig. 6a, right). Finally, lengths of microtubules were estimated by projecting the locations of the EbA-mCherry puncta along the traced hyphal axis (yellow line in Fig. 6a).

**FIG. 6:**
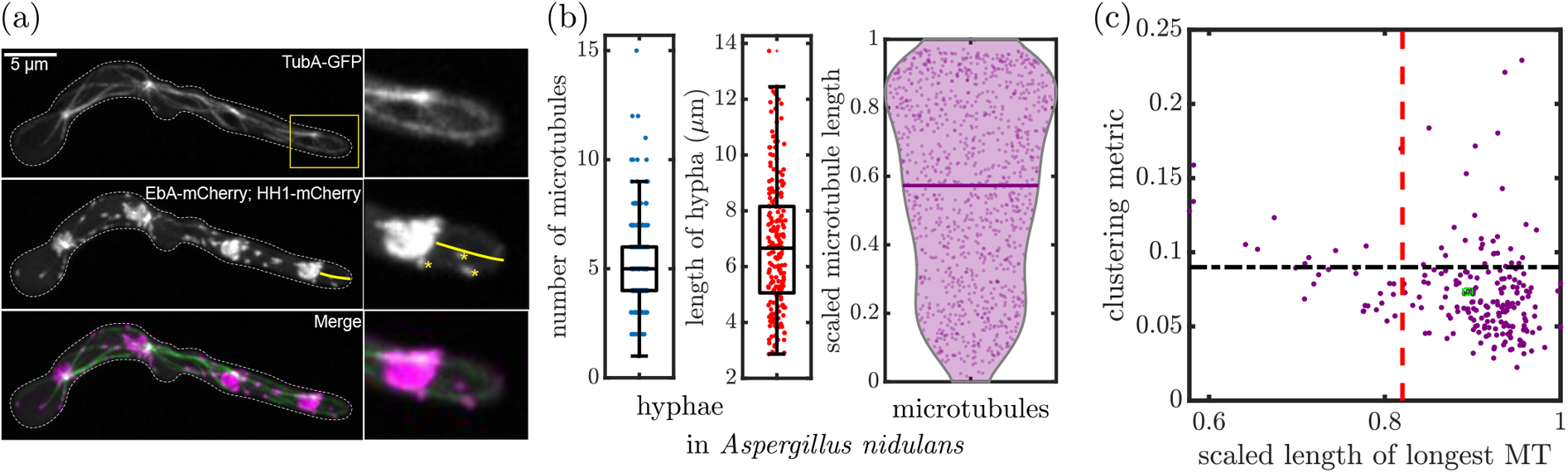
Microtubule configurations in *Aspergillus nidulans* hyphae. (a) Micrographs of *A. nidulans* germlings expressing fluorescently tagged tubulin (TubA-GFP, top), plus-end binding protein EbA/EB1 (EbA-mCherry), and nuclei (HH1-mCherry, middle panel). White dotted line shows the outline of the hypha. Yellow box denotes the cropped region shown on the right. Yellow line denotes measured length between hyphal tip and closest nucleus. Yellow asterisks denote EB1 plus-ends. (b) Number of microtubules (left), hypha length (center), and scaled microtubule length (right) for *n* = 210 hyphal tip regions. Microtubule lengths are scaled by the length of the corresponding region from the last nucleus to the cell tip. (c) Scatter plot showing the scaled length of the longest microtubule, and the clustering metric for hyphal microtubule configurations. The mean value for these metrics is indicated by the green point (scaled longest MT 0.89 *±* 0.005; clustering metric *d* = 0.073 *±* 0.002). The vertical red line (0.82 *±* 0.0002) and the horizontal black line (0.09 *±* 0.00004) denote the average value of each corresponding metric for 10^6^ uniformly sampled configurations. All intervals and error bars correspond to mean *±* standard error.

Based on data from *n* = 210 hyphae, the average length of the region from the last nucleus to the tip was 0.16*μ*m (s.e.m.). Each post-nuclear hypha region contained 5.13 ± 0.15 (s.e.m.) microtubules with an average length of 4.15±0.08*μ*m (s.e.m.). Fig. 6b shows the distributions of the observed hypha and microtubule lengths. Due to the large variability in cell size, we scale microtubule lengths with respect to the length of the individual hypha. For all hyphae, scaled length of the longest microtubule and the minimal-distance clustering metric for microtubule plus-end positions is calculated and plotted in Fig. 6c.

We compare microtubule arrangements in *A. nidulans* to the null hypothesis of microtubule ends scattered uniformly throughout the domain. This comparison helps identify non-uniform features of the microtubule distribution which can then be compared to our computational predictions for optimal arrangements. To this end, we generate 10^6^ randomly sampled microtubule configurations. Each configuration has a number of microtubules drawn from the distribution observed in *A. nidulans* hyphae (Fig. 6b, left), with each scaled microtubule length selected uniformly at random (between 0 and 1). The longest microtubule length and the clustering metric is computed for each random configuration, and the average values are plotted as dashed lines on Fig. 6c.

The quantification of observed hyphal microtubule configurations demonstrated substantial differences from the null hypothesis of uniformly distributed random configurations. Namely, the mean scaled length of the longest microtubule was significantly longer than the value that would be expected for uniform architectures (*p* < 0.001 from one-sided t-test). Furthermore, the mean clustering metric for *A. nidulans* microtubules (0.073 ± 0.002) is significantly lower than the 0.09 ± 0.00004 value for random configurations (*p* < 0.001 from one-sided t-test). It should be noted, by contrast, that configurations selected specifically for long microtubules would be expected to have a clustering metric substantially above the uniformly distributed value, due to accumulation of multiple plus-ends at the distal tip. These comparisons indicate that microtubule arrangements in *A. nidulans* hyphae tend to have one long microtubule, with the remaining plus-ends broadly distributed throughout the domain. These features match the optimal configuration predicted from the computational model for particle capture at plus-ends. The hyphal measurements highlight the fact that microtubules exhibit a distinctly non-uniform, yet non-clustered, length distribution that should lead to efficient capture of both particles entering at the cell tip and those produced throughout the entire hypha.

## IV. CONCLUSIONS

We have employed analytical modeling and computational simulations to highlight the role of microtubule arrangements in capturing cargo within tubular cells. For cargo entering at the distal end of the cell and captured at microtubule plus-ends, we show that spreading capture regions away from the entry point results in faster engagement with microtubules. The effect of cell size on such optimal arrangements is explored, revealing that for cell lengths on the order of 10*μ*m, it is optimal to distribute microtubule ends over up to 25% of the axial length. In longer cells, it becomes advantageous to cluster the plus-ends over a relatively smaller fraction of the domain. Some cargos require a period of maturation before becoming available to engage with microtubules, and we show that maturation processes on time-scales above a few minutes require a broader distribution of microtubule ends for efficient capture.

By analyzing random microtubule configurations, we establish general principles for rapid cargo capture across various scenarios for initial cargo distribution and length of the capture regions. We show that configurations with a single long microtubule reaching the cell tip, accompanied by broad dispersal of the remaining microtubule ends, are ideal for rapidly capturing a variety of cargo. Such distributions can be established by tuning microtubule catastrophe rates as highlighted by simulations of cargo capture in a minimal model of dynamic microtubules. Finally, we image microtubules in *A. nidulans* hyphae and show that their length distributions follow the general principles for optimality laid down by our model. These results highlight novel aspects of cytoskeletal organization and its impact or cargo capture, providing possible mechanisms to establish optimal arrangements and validating predictions using *in vivo* data.

A central challenge for microtubule organization in a cell is the necessity for a single cytoskeletal architecture to serve as a transport highway for a variety of different cargos with different transport objectives. In this study, we focused specifically on the retrograde delivery of cargo to the nuclear region. However, other cargos require delivery from the nucleus to the periphery or broad distribution throughout the cellular domain. These alternate objectives impose different utility functions on the possible microtubule configurations. For directed delivery, long microtubules enable cargo to be deposited close to the distal tip, but axial separation of microtubule end positions has been shown to promote retention at the tip by reducing the recirculation of entrained cytoplasmic fluid [46]. For broad distribution of bidirectionally moving cargo, short microtubules may reduce the processive run-time of directed transport [19], resulting in increased frequency of reversals and more rapid distribution of cargo [12]. Architectures with short microtubules of mixed polarity can also enable dispersion of cargos whose motion is dominated by a single motor type, as observed in the proximal regions of mammalian dendrites [47]. The analytically tractable 1D modeling approach and 3D simulations developed here can be extended in future work to consider the impact of microtubule length distributions on this broad variety of intracellular transport systems.

By delineating the role of microtubule arrangements in individual transport processes, we can begin to gain a comprehensive picture of the evolutionary pressures guiding the observed microtubule architectures in live cells. Furthermore, establishing the impact of cytoskeletal morphology on the efficiency of key transport objectives is critical to developing a predictive understanding of how pharmacological or genetic perturbations in cytoskeletal filament length modulate cellular functions.

## Acknowledgements

This work was funded by a National Science Foundation CAREER Award (1848057). J.R.C. is funded by a MOSAIC K99/R00 award from the National Institutes of Health (K99GM140269), and S.L.R.-P. is funded by the Howard Hughes Medical Institute and the National Institutes of Health (R01GM121772). We thank the Nikon Imaging Center at UC San Diego.

## References

[1] M. Burute and L. C. Kapitein, Annu Rev Cell Dev Biol 35, 29 (2019).

[2] S. S. Mogre, A. I. Brown, and E. F. Koslover, Phys Biol 17, 061003 (2020).

[3] A. D. Sanchez and J. L. Feldman, Curr Opin Cell Biol 44, 93 (2017).

[4] P. W. Baas and F. J. Ahmad, Brain 136, 2937 (2013).

[5] D. Ando, N. Korabel, K. C. Huang, and A. Gopinathan, Biophys J 109, 1574 (2015).

[6] P. J. Mlynarczyk and S. M. Abel, Phys Rev E 99, 022406 (2019).

[7] M. Scholz, S. Burov, K. L. Weirich, B. J. Scholz, S. A. Tabei, M. L. Gardel, and A. R. Dinner, Phys Rev X 6, 011037 (2016).

[8] M. Scholz, K. L. Weirich, M. L. Gardel, and A. R. Dinner, Soft Matter 16, 2135 (2020).

[9] A. T. Lombardo, S. R. Nelson, G. G. Kennedy, K. M. Trybus, S. Walcott, and D. M. Warshaw, P Natl Acad Sci 116, 8326 (2019).

[10] A. E. Hafner and H. Rieger, Biophys J 114, 1420 (2018).

[11] B. Maelfeyt, S. A. Tabei, and A. Gopinathan, Phys Rev E 99, 062404 (2019).

[12] S. S. Mogre and E. F. Koslover, Phys Rev E 97, 042402 (2018).

[13] C. F. Ibáñez, Trends Cell Biol 17, 519 (2007).

[14] P. D. Chowdary, D. L. Che, and B. Cui, Annu Rev Phys Chem 63, 571 (2012).

[15] S. Maday and E. L. Holzbaur, Autophagy 8, 858 (2012).

[16] C. Barlowe and A. Helenius, Annu Rev Cell Dev Biol 32, 197 (2016).

[17] L. C. Kapitein and C. C. Hoogenraad, Mol Cell Neurosci 46, 9 (2011).

[18] M. J. Egan, M. A. McClintock, and S. L. Reck-Peterson, Curr Opin Microbiol 15, 637 (2012).

[19] S. Yogev, R. Cooper, R. Fetter, M. Horowitz, and K. Shen, Neuron 92, 449 (2016).

[20] G. Han, B. Liu, J. Zhang, W. Zuo, N. R. Morris, and X. Xiang, Curr Biol 11, 719 (2001).

[21] M. Schuster, S. Kilaru, P. Ashwin, C. Lin, N. J. Severs, and G. Steinberg, Embo J 30, 652 (2011).

[22] A. J. Moughamian, G. E. Osborn, J. E. Lazarus, S. Maday, and E. L. Holzbaur, J Neurosci 33, 13190 (2013).

[23] K. T. Vaughan, S. H. Tynan, N. E. Faulkner, C. J. Echeverri, and R. B. Vallee, J Cell Sci 112, 1437 (1999).

[24] J.-H. Lenz, I. Schuchardt, A. Straube, and G. Steinberg, Embo J 25, 2275 (2006).

[25] J. A. Perge, J. E. Niven, E. Mugnaini, V. Balasubramanian, and P. Sterling, J Neurosci 32, 626 (2012).

[26] S. S. Mogre, J. R. Christensen, C. S. Niman, S. L. Reck-Peterson, and E. F. Koslover, Biophys J 118, 1357 (2020).

[27] C. Lin, M. Schuster, S. C. Guimaraes, P. Ashwin, M. Schrader, J. Metz, C. Hacker, S. J. Gurr, and G. Steinberg, Nat Commun 7, 11814 (2016).

[28] S. Redner, A guide to first-passage processes (Cambridge University Press, 2001).

[29] Z. C. Scott, A. I. Brown, S. S. Mogre, L. M. Westrate, and E. F. Koslover, arXiv preprint arXiv:2103.05065 (2021).

[30] G. Steinberg, R. Wedlich-Soldner, M. Brill, and I. Schulz, J Cell Sci 114, 609 (2001).

[31] R. Pordes, D. Petravick, B. Kramer, D. Olson, M. Livny, A. Roy, P. Avery, K. Blackburn, T. Wenaus, F. Würthwein, et al., in J. Phys. Conf. Ser. (2007), vol. 78 of 78, p. 012057.

[32] I. Sfiligoi, D. C. Bradley, B. Holzman, P. Mhashilkar, S. Padhi, and F. Wurthwein, in 2009 WRI World Congress on Computer Science and Information Engineering (2009), vol. 2 of 2, pp. 428–432.

[33] D. H. Smith, Prog Neurobiol 89, 231 (2009).

[34] X.-T. Cheng, B. Zhou, M.-Y. Lin, Q. Cai, and Z.-H. Sheng, J Cell Biol 209, 377 (2015).

[35] Z. Xie and D. J. Klionsky, Nat Cell Biol 9, 1102 (2007).

[36] S. Maday, K. E. Wallace, and E. L. Holzbaur, J Cell Biol 196, 407 (2012).

[37] Z. S. Schultzhaus and B. D. Shaw, Fungal Biology Reviews 29, 43 (2015).

[38] L. C. Kapitein and C. C. Hoogenraad, Neuron 87, 492 (2015).

[39] C. Conde and A. Cáceres, Nat Rev Neurosci 10, 319 (2009).

[40] J. Salogiannis and S. L. Reck-Peterson, Trends Cell Biol 27, 141 (2017).

[41] J. Salogiannis, M. J. Egan, and S. L. Reck-Peterson, J Cell Biol 212, 289 (2016).

[42] K. Tan, A. J. Roberts, M. Chonofsky, M. J. Egan, and S. L. Reck-Peterson, Mol Biol Cell 25, 669 (2014).

[43] J. F. Abenza, A. Galindo, M. Pinar, A. Pantazopoulou, V. de los Ríos, and M. A. Peñalva, Mol Biol Cell 23, 1889 (2012).

[44] G. Steinberg, Curr Opin Microbiol 20, 10 (2014).

[45] M. Bartoszewska, Ł. Opaliński, M. Veenhuis, and I. J. van der Klei, Biotechnol Lett 33, 1921 (2011).

[46] P. K. Trong, J. Guck, and R. E. Goldstein, Phys Rev Lett 109, 028104 (2012).

[47] L. C. Kapitein, M. A. Schlager, M. Kuijpers, P. S. Wulf, M. van Spronsen, F. C. MacKintosh, and C. C. Hoogenraad, Curr Biol 20, 290 (2010).

