## Supplemental material for "Optimizing microtubule arrangements for rapid cargo capture"

#### S1. METHODS FOR GROWING AND IMAGING *ASPERGILLUS NIDULANS* STRAINS USED IN THIS STUDY

*Aspergillus nidulans* strains were grown on yeast extract and glucose media agar gum plates for maintenance [1]. For spinning disk microscopy of *A. nidulans* germlings, *A. nidulans* spores were resuspended in 1 mL of 0.01% Tween-80. The spore/Tween-80 solution was then added 1:1000 to 1% glucose minimal media with no supplements in a 4-chamber 35mm dish with #1.5 coverglass bottom (Cellvis), and incubated for 16-20 hours at 30°C. Germlings were imaged using a Yokogawa W1 confocal scanhead mounted to a Nikon Ti2 microscope with an Apo TIRF 100x 1.49 NA objective. The scope was run with NIS Elements using the 488nm and 561nm lines of a six-line (405nm, 445nm, 488nm, 515nm, 561nm, and 640nm) LUN-F-XL laser engine and a Prime95B camera (Photometrics). Image channels were acquired sequentially using bandpass filters for each channel (525/50 and 595/50). Z-stacks were acquired using a piezo Z stage (Mad City Labs). The z-range used to image a field of germlings was set manually depending on germling extension from the coverglass surface.

The *Aspergillus nidulans* strain used in this study is listed in Table I. Strain RPA361 expressing EbA-mCherry, TubA-GFP, and HH1-mCherry was created through genetic crossing, as previously described [2].

| Strain | Genotype | Source |
|---|---|---|
| RPA361 | *[ebA*-mCherry-*Afribo]*, *[tubA*-GFP-*Afpyro]*; *[HH1*-mCherry-*AfPyrG]*; *riboB2*; *pyroA4*; *pyrG89*; $\Delta$*nkuA::bar* | This study |

Table I: *A. nidulans* strain used in this study

### S2. EFFECT OF CAPTURE REGION SIZE ON CARGO CAPTURE TIME

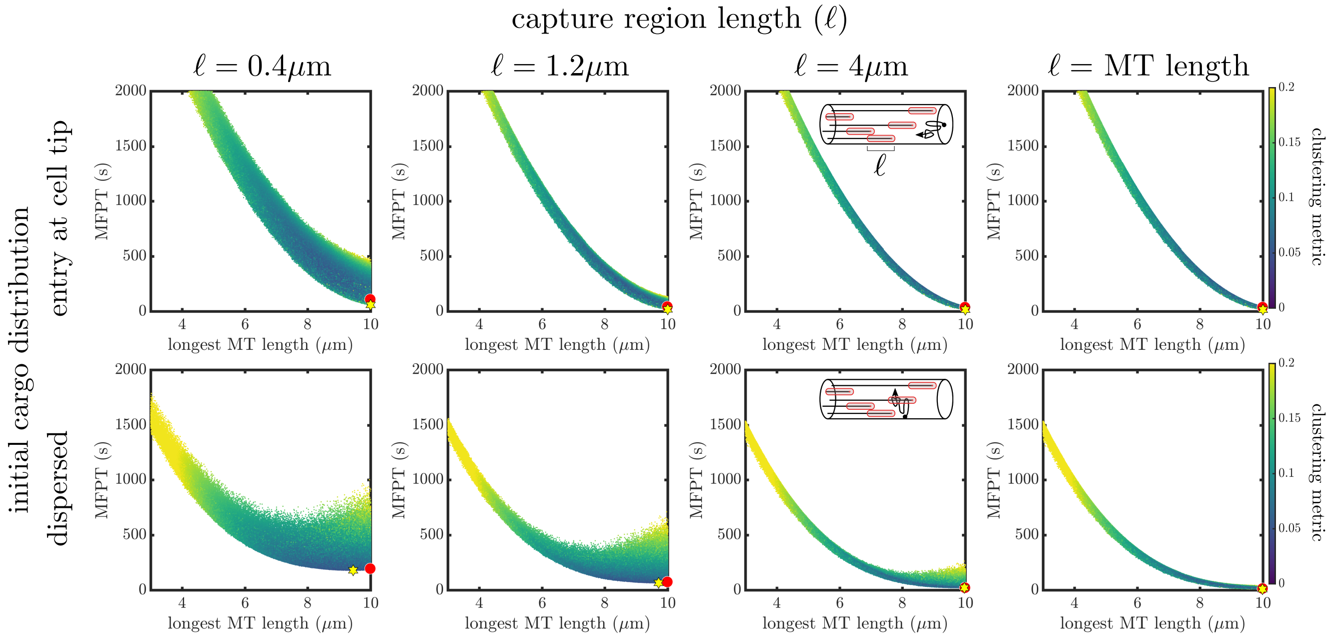


Figure S1: **Cargo capture by regions of varying size.** Scatter plots showing the MFPT vs the length of the longest microtubule for $10^6$ randomly sampled configurations with 5 microtubules each in a domain of length $10\mu$m. The length of the capture region is indicated by $\ell$. The top row denotes the MFPT for cargo formed at the cell tip. The bottom row denotes the MFPT for cargo initially dispersed uniformly. The red circle denotes a configuration that falls within the lowest 2.5% of MFPTs for both starting distributions for capture near plus-ends ($\ell = 0.4\mu$m). Yellow stars denote the overall fastest configuration for a given capture condition. The color of the scatter points denotes the clustering metric described in the main text.

In the main text, we focused on cargo loaded onto microtubules only within a 200nm contact radius of the plus-end. However, dynein comets generally exhibit a gradual decrease in density over a micrometer length scale [3]. Furthermore, some cargos may recruit their own motor protein assembly and should be able to initiate retrograde transport elsewhere along the microtubule. For a given arrangement of microtubule lengths, the cargo capture time can vary widely depending on the size and availability of regions along the microtubule where the cargo can bind. In this section, we quantify the relation between the size of the capture region along a microtubule and the MFPT to capture cargo.

Fig. S1 shows the MFPT for $10^6$ randomly sampled configurations of 5 microtubules, for several different values of the capture region length $\ell$ (equivalent to twice the capture radius). The left-most plots correspond to Fig. 4a. The right-most plots represent the limiting case where cargo can be captured along the entire microtubule. In this limiting case, we see that the longest-microtubule length is a strong predictor of the capture time, with very little variation among the MFPT for different configurations with the same longest length. Intermediate values of the capture length behave essentially as an interpolation between narrow capture at the tip and capture along the whole microtubule.

The red dots in Fig. S1 correspond to the configuration shown in Fig. 4b.iii, which performs nearly optimally for plus-end capture with both cargo entering at the cell tip and cargo starting with a uniform distribution. This configuration has a single microtubule stretching all the way to the distal tip, with the other microtubule ends spaced out evenly throughout the domain. Notably, such a configuration also falls within the lowest 3% of MFPTs for the case with capture along the entire microtubule. Thus microtubule architectures with these dual features are near-optimal for rapid initiation of retrograde transport in a broad variety of scenarios, including different cargo entry points and different lengths of microtubule capture regions.

[1] E. Szewczyk, T. Nayak, C. E. Oakley, H. Edgerton, Y. Xiong, N. Taheri-Talesh, S. A. Osmani, and B. R. Oakley, Nat Protoc **1**, 3111 (2006).

[2] R. B. Todd, M. A. Davis, and M. J. Hynes, Nat Protoc **2**, 811 (2007).

[3] M. Schuster, S. Kilaru, P. Ashwin, C. Lin, N. J. Severs, and G. Steinberg, Embo J **30**, 652 (2011).
